# Crosstalk in skin: Loss of desmoglein 1 in keratinocytes inhibits BRAF^V600E^-induced cellular senescence in human melanocytes

**DOI:** 10.1101/2023.02.16.528886

**Authors:** Xin Tong, Hope E. Burks, Ziyou Ren, Jennifer L. Koetsier, Quinn R. Roth-Carter, Kathleen J. Green

## Abstract

Melanoma arises from transformation of melanocytes in the basal layer of the epidermis where they are surrounded by keratinocytes, with which they interact through cell contact and paracrine communication. Considerable effort has been devoted to determining how the accumulation of oncogene and tumor suppressor gene mutations in melanocytes drive melanoma development. However, the extent to which alterations in keratinocytes that occur in the developing tumor niche serve as extrinsic drivers of melanoma initiation and progression is poorly understood. We recently identified the keratinocyte-specific cadherin, desmoglein 1 (Dsg1), as an important mediator of keratinocyte:melanoma cell crosstalk, demonstrating that its chronic loss, which can occur through melanoma cell-dependent paracrine signaling, promotes behaviors that mimic a malignant phenotype. Here we address the extent to which Dsg1 loss affects early steps in melanomagenesis. RNA-Seq analysis revealed that paracrine signals from Dsg1-deficient keratinocytes mediate a transcriptional switch from a differentiated to undifferentiated cell state in melanocytes expressing BRAF^V600E^, a driver mutation commonly present in both melanoma and benign nevi and reported to cause growth arrest and oncogene-induced senescence (OIS). Of ∼220 differentially expressed genes in BRAF^V600E^ cells treated with Dsg1-deficient conditioned media (CM), the laminin superfamily member NTN4/Netrin-4, which inhibits senescence in endothelial cells, stood out. Indeed, while BRAF^V600E^ melanocytes treated with Dsg1-deficient CM showed signs of senescence bypass as assessed by increased senescence-associated β-galactosidase activity and decreased p16, knockdown of NTN4 reversed these effects. These results suggest that Dsg1 loss in keratinocytes provides an extrinsic signal to push melanocytes towards oncogenic transformation once an initial mutation has been introduced.

## Introduction

Cutaneous melanoma is an aggressive skin cancer arising from the pigment-producing melanocytes of the human epidermis, and its incidence is increasing worldwide [1]. While the presence of driver mutations that promote melanoma initiation and development is well-established, their presence does not guarantee that a melanoma will develop, nor do all melanomas arise from nevi expressing driver mutations [2, 3]. Analysis of matched primary and metastatic melanomas have failed to identify definitive genetic/mutational signatures that dictate melanoma progression [4, 5]. On the other hand, it is also appreciated that extrinsic factors emanating from the tumor environment can drive transcriptional changes in tumor cells that lead not only to tumor progression but also the acquisition of drug resistance [6]. Thus, understanding what external factors cooperate with genetic changes in melanoma cells will be critical for establishing mechanisms of tumor progression.

Activating mutations in BRAF, a gene encoding for a serine/threonine protein kinase that drives growth signals, are commonly found in melanoma tumors. Over half of all cutaneous melanoma harbor the BRAF^V600E^ mutation, which leads to constitutive activation of the mitogen-activated protein kinase (MAPK) pathway [7-9]. After an initial burst of proliferation, melanocytes that acquire the BRAF^V600E^ mutation enter a state of growth arrest, which has been attributed to oncogene-induced senescence (OIS) [10]. Thus, while BRAF^V600E^ is present in up to 80% of benign nevi [7], these nevi rarely progress toward melanoma. While melanocytes within these preneoplastic lesions are positive for multiple senescence markers [7], alternative mechanisms by which nevus cells might be growth arrested have recently been proposed [11, 12]. Regardless, escaping this growth arrested state is an important step in melanoma development. Studies characterizing senescence and growth arrest bypass have focused primarily on those emanating from loss of function mutations in tumor suppressor genes such as the Wnt signaling effector *β*-catenin, the tumor suppressor PTEN, or cell cycle regulators like CDKN2A/B [13-16]. While the importance of extrinsic factors driving melanoma progression is now appreciated, less attention has been paid to non-genetic factors that might prevent or reverse a growth-arrested state in early melanomagenesis.

One potential source of such extrinsic signals is the keratinocytes, which surround epidermal melanocytes and outnumber them ∼36:1. Keratinocytes secrete factors such as melanocyte-stimulating hormones that facilitate the synthesis of melanin in melanocytes, which is then packaged and sent back to keratinocytes to protect them from ultraviolet radiation (UV)-mediated DNA damage [17-20]. In the skin, we recently identified the keratinocyte-specific cadherin Desmoglein 1 (Dsg1) as a key factor of communication between keratinocytes and melanocytes [21]. Best known as a cell-cell junction protein unique to stratified tissues of terrestrial vertebrates [22], Dsg1 has distinctive properties that position it to be a sensor of environmental stress beyond its role as an adhesion molecule [23, 24]. While transient Dsg1 downregulation like that occurring in response to UV initiates a protective response in melanocytes [21, 25], chronic loss stimulates pathogenic pro-inflammatory cytokine and chemokine production [26], and promotes melanocyte behaviors associated with transformation, such as pagetoid movement in 3D organotypic cultures [21]. We also recently demonstrated that conditioned media from Dsg1-deficient keratinocytes can increase migration in melanoma cells through CXCL1/CXCR2 signaling axis [27]. However, a potential role for the paracrine signaling caused by keratinocyte Dsg1 loss in the early steps of melanocyte transformation is unknown.

In the present study, we introduced wild-type (WT) BRAF and the constitutively active BRAF^V600E^ into human primary melanocytes and addressed the role of paracrine signaling from Dsg1-deficient keratinocytes through conditioned media (CM) treatment. RNA-Seq analysis revealed that paracrine signals from Dsg1-deficient keratinocytes mediated a transcriptional switch from a differentiated to undifferentiated cell state, only in BRAF^V600E^ expressing cells, whereas increased stem-like and invasive signatures occurred in control (untransduced) melanocytes, WT BRAF and BRAF^V600E^ expressing cells. As expected, we observed an increase in senescence-associated β-galactosidase (SA-β-Gal) activity and elevated p16 protein levels in BRAF^V600E^-transduced melanocytes [28, 29]. Importantly, CM from Dsg1-deficient keratinocytes reversed signs of BRAF^V600E^-induced senescence in a portion of transduced cells. Among up-regulated genes in BRAF^V600E^-transduced melanocytes were several adhesion and matrix molecules, including NTN4/Netrin-4, a cellular senescence inhibitor [30-32]. Knockdown of NTN4 reversed senescence bypass mediated by Dsg1-deficient CM, supporting its role in bypassing BRAF^V600E^-induced senescence. Collectively, our results suggest that Dsg1 loss in keratinocytes provides a non-genetic, extrinsic signal capable of promoting oncogenic transformation in melanocytes expressing a single driver mutation.

## Results

### Transcriptomic analysis reveals a transcriptional switch in BRAF^V600E^ melanocytes induced by loss of keratinocyte Dsg1

Recently, we reported evidence of a feed forward pro-melanomagenic loop in which melanoma cells reduce Dsg1 expression in keratinocytes, and, in turn, paracrine signaling from Dsg1-deficient keratinocytes promotes melanoma cell migration [27]. How paracrine signaling from Dsg1-deficient keratinocytes might act earlier in melanocyte transformation was not tested. Given the importance of extrinsic factors in promoting tumor progression, and since most melanocytes with driver mutations (like BRAF mutations in nevi) rarely go on to produce melanoma, we asked whether the loss of Dsg1 in keratinocytes can promote further steps in melanocyte transformation.

Toward this end, we addressed the role of paracrine signaling from Dsg1-deficient keratinocytes to primary melanocytes expressing mutant BRAF, as this gene is mutated in up to 80% of benign nevi, where it drives OIS/growth arrest to suppress uncontrolled proliferation of melanocytes [7, 10]. Approximately 80–90% of BRAF mutations are V600E (valine to glutamic acid) missense mutation [33, 34]. Therefore, we introduced BRAF^V600E^ or wild type (WT) BRAF into human primary melanocytes through lentivirus transduction to build an in vitro BRAF^V600E^-induced senescence model. Validating this model, we saw an increase in senescence-associated β-galactosidase (SA-β-Gal) activity, a hallmark of senescence, by X-gal staining in BRAF^V600E^-but not WT BRAF-transduced melanocytes (Fig. S1A and S1B). Similar results were obtained with a more sensitive flow cytometry assay for SA-β-Gal activity (Fig. S1C and S1D). Elevated expression of p16, another indicator of senescence, was also induced by BRAF^V600E^ but not WT BRAF as expected (Fig. S1E).

Having validated this model system, we set out to establish transcriptional changes in control (untransduced) primary melanocytes, and melanocytes expressing WT BRAF or BRAF^V600E^. CM was collected from each cell type into which shDsg1 (or shCTL) was delivered using retroviral transduction to knock down Dsg1 in keratinocytes (Fig. S2A)). Each of the 6 experimental arms was performed with three matched keratinocyte:melanocyte pairs. After batch correction to adjust for the pair effects, we performed gene set enrichment analysis (GSEA) on control, WT BRAF and BRAF^V600E^ transduced melanocytes treated with Dsg1 knockdown or control keratinocyte CM using previously published melanoma and melanocyte gene signatures [35-42]. Pathway analysis revealed that in all melanocyte conditions, treatment with CM from Dsg1 knockdown keratinocytes led to a general enrichment in pathways associated with a migratory phenotype in all genetic backgrounds--control, WT BRAF and BRAF^V600E^ transduced cells (Fig. 1A). However, alterations in gene signatures associated with pigmentation and melanocytic differentiation status differed in control and WT BRAF cells compared with BRAF^V600E^ transduced cells. Pathways associated with pigmentation and differentiation were markedly upregulated in control and WT BRAF-transduced melanocytes upon treatment with Dsg1-deficient CM. In contrast, those same pathways were significantly downregulated in melanocytes expressing the exogenous BRAF^V600E^ (Fig. 1A). These data are interesting considering our previously published work showing that Dsg1-deficient keratinocytes promote pagetoid movement of melanocytes in 3D co-cultures, and increased pigmentation secretion when treated with Dsg1-deficient media [21].

**Figure 1.**
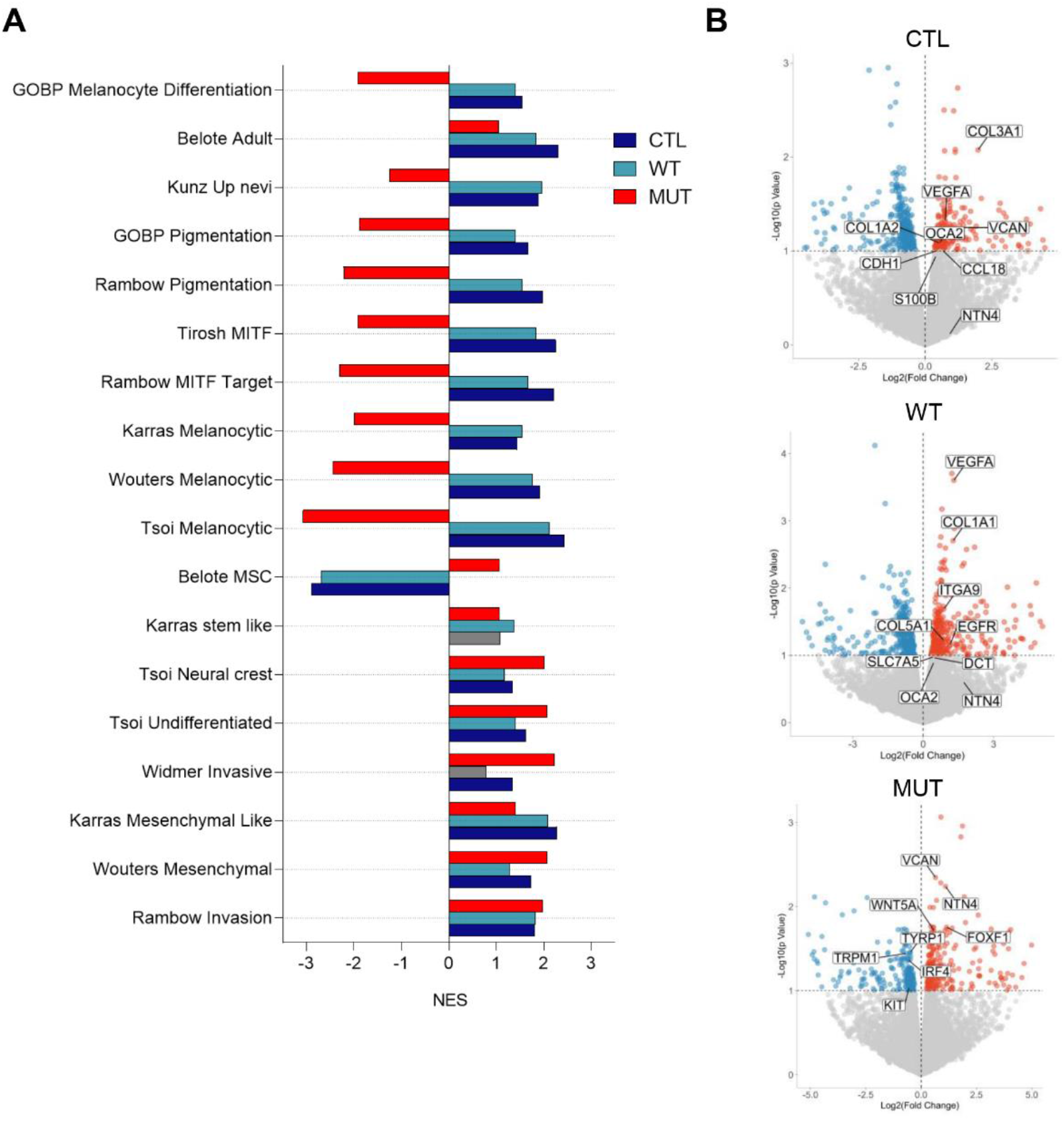
Loss of Dsg1 in keratinocytes induces oncogenic transformation of BRAF^V600E^-transduced melanocytes. **A**, Gene set enrichment analysis compared differentially expressed gene sets to published signatures of melanoma/melanocyte signaling. NES, normalized enrichment score. **B**, Volcano plots revealed log2 fold change against nominal p values (-log10) from differential expression analysis of mRNA sequencing of melanocytes with different BRAF statuses (control, WT BRAF and BRAF^V600E^) and treated with control or Dsg1 knockdown conditioned media. Select genes involved in differentiation, pigmentation, and proliferation are marked.

Differential gene expression analysis comparing control, WT BRAF and BRAF^V600E^ transduced melanocytes treated with Dsg1 knockdown or control keratinocyte CM revealed ∼170, 210 and 220 genes that were significantly differentially expressed, respectively (nominal p value<0.05) (Fig. 1B). Among them, we found collagen genes such as COL1A1, COL3A1, COL5A1 (invasion/motility), VEGFA (proliferation/migration), and OCA2 (melanin synthesis), were upregulated in control and WT BRAF transduced melanocytes treated with Dsg1-deficient keratinocyte CM. On the other hand, TYRP1, TRPM1, IRF4 and KIT (pigmentation/differentiation) were downregulated and FOXF1 (melanoma cell proliferation) was upregulated in BRAF^V600E^ transduced melanocytes treated with Dsg1-deficient keratinocyte CM (Fig. 1B). Genes involved in cell adhesion and matrix biology were also enriched in BRAF^V600E^ as discussed in more detail below.

### Conditioned media from Dsg1-deficient keratinocytes inhibits BRAF^V600E^-induced senescence in melanocytes

Given the transcriptional signatures uniquely arising after different CM treatment, we set out to determine if Dsg1-deficient environment also drove functional changes in these cells. Compared to control melanocytes and melanocytes transduced with WT BRAF, which continued to grow in CM (either shCTL or shDsg1 CM), we found that BRAF^V600E^-transduced melanocytes stopped growing soon after BRAF^V600E^ transduction. After 3 days of CM treatment, the number of BRAF^V600E^-transduced melanocytes was dramatically less than those of control/WT BRAF-transduced melanocytes (Fig. 2A).

**Figure 2.**
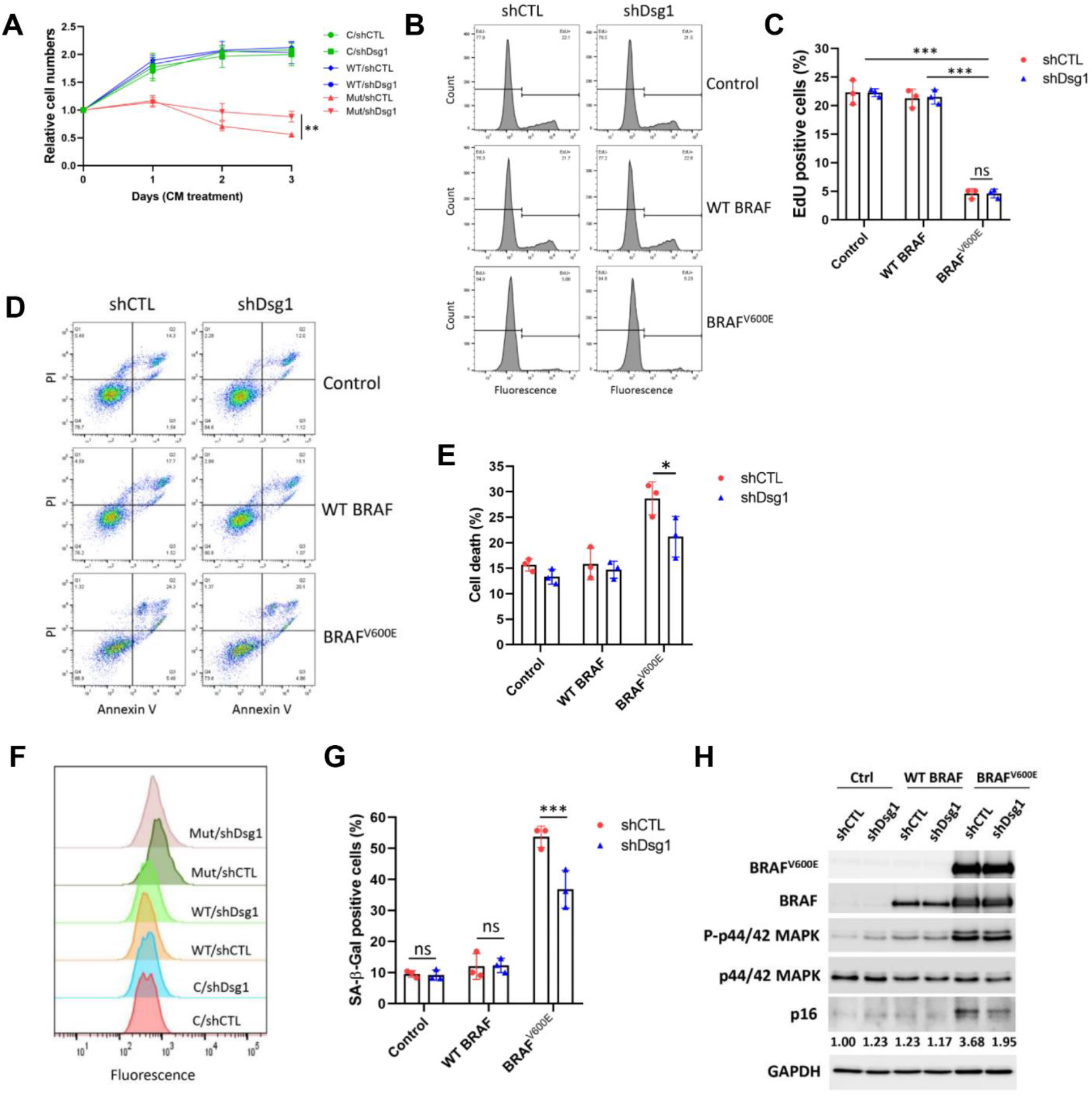
Dsg1-deficient conditioned media (CM) suppresses BRAF^V600E^-induced senescence in melanocytes. Human primary melanocytes were transduced with WT BRAF or BRAF^V600E^ lentivirus, control or Dsg1-deficient CM were added 24 h after transduction and cells continued to be cultured in CM before they were harvested. **A**, Relative cell numbers over time. **B**, EdU flow cytometry assay to determine cell proliferation. The figures showed a clear separation of proliferating cells which have incorporated EdU in control and WT BRAF melanocytes but not in BRAF^V600E^ melanocytes. **C**, Quantification of EdU flow cytometry assay. **D**, Annexin V/dead cell apoptosis flow cytometry assay was performed to detect cell death. **E**, Quantification of cell death by flow cytometry assay. **F**, Senescence-associated β-galactosidase activity was detected by flow cytometry assay. Right shift of the peak indicated the increase in senescence. **G**, Quantification of senescence flow cytometry assay. **H**, Expression of the indicated proteins was determined by Western blot. Results were reported as mean ± SD (n=3), statistical analysis was performed using two-way ANOVA with multiple comparisons (*, P < 0.05; ***, P < 0.001).

Interestingly, total cell number was greater in BRAF^V600E^-transduced melanocyte populations treated with Dsg1-deficient CM than those treated with control CM (Fig. 2A). To address whether this difference was due to elevated cell proliferation in cells treated with Dsg1 knockdown CM, an EdU incorporation experiment was carried out. While EdU positive cells were substantially less in BRAF^V600E^-transduced melanocytes than in control and WT BRAF-transduced melanocytes, no difference in EdU incorporation between the two CM-treated cell populations was observed (Fig. 2B and 2C). To determine whether difference in cell death underlie the observed difference in cell numbers after the two CM treatments, we assessed cell death (both apoptosis and necrosis) by flow cytometry. As illustrated in Fig. 2D, when both types of cell death were monitored (apoptosis plus necrosis, windows Q2 plus Q3 in Fig. 2D), we found that there was significantly less death of cells treated with Dsg1-deficient CM (Fig. 2E).

Having established that BRAF^V600E^ expressing melanocytes exhibit signs of OIS, we next assessed the extent to which shDsg1 CM media can reverse OIS-related features in these cells. The shDsg1 CM treated BRAF^V600^ melanocytes exhibited a partial but significant decrease in SA-β-Gal activity as demonstrated by the flow cytometry senescence assay (Fig 2F and 2G). X-gal staining showed a similar result (Fig. S2B and S2C)). We also observed a reduction in the protein product of CDKN2A, p16, expression in BRAF^V600E^ melanocytes treated with shDsg1 CM compared to control CM (Fig. 2H). Together, these results demonstrated that CM from Dsg1-deficient keratinocytes results in senescence bypass in a reproducible proportion of BRAF^V600E^ expressing melanocytes.

### Netrin-4 plays an important role in Dsg1 loss-induced senescence bypass

To investigate the underlying mechanisms responsible for the bypass of senescence and increased cell numbers induced by Dsg1-deficient CM, we returned to the RNA-seq data of CM treated BRAF^V600E^-transduced samples. Gene Ontology (GO) pathway analysis revealed that gene sets up-regulated by Dsg1-deficient CM included signatures associated with cell adhesion molecules, cell-cell adhesion and extracellular matrix organization, in addition to more specific matrix-associated pathways like signaling through the neuronal guidance molecule Netrin-1 (Fig. 3A). To gain a further understanding of specific genes that could contribute to Dsg1 loss-induced senescence bypass we performed cluster analysis (Fig. 3B). Among the top 50 changed genes, the laminin superfamily gene and Netrin-1 relative NTN4/Netrin-4 was upregulated in BRAF^V600E^-transduced melanocytes. Netrin-4 has recently been reported as a cellular senescence inhibitor, preventing endothelial senescence and DNA damage-induced senescence in glioblastoma [30-32]. Therefore, we considered it as a good candidate for mediating senescence bypass in BRAF^V600E^ melanocytes treated with shDsg1 CM.

**Figure 3.**
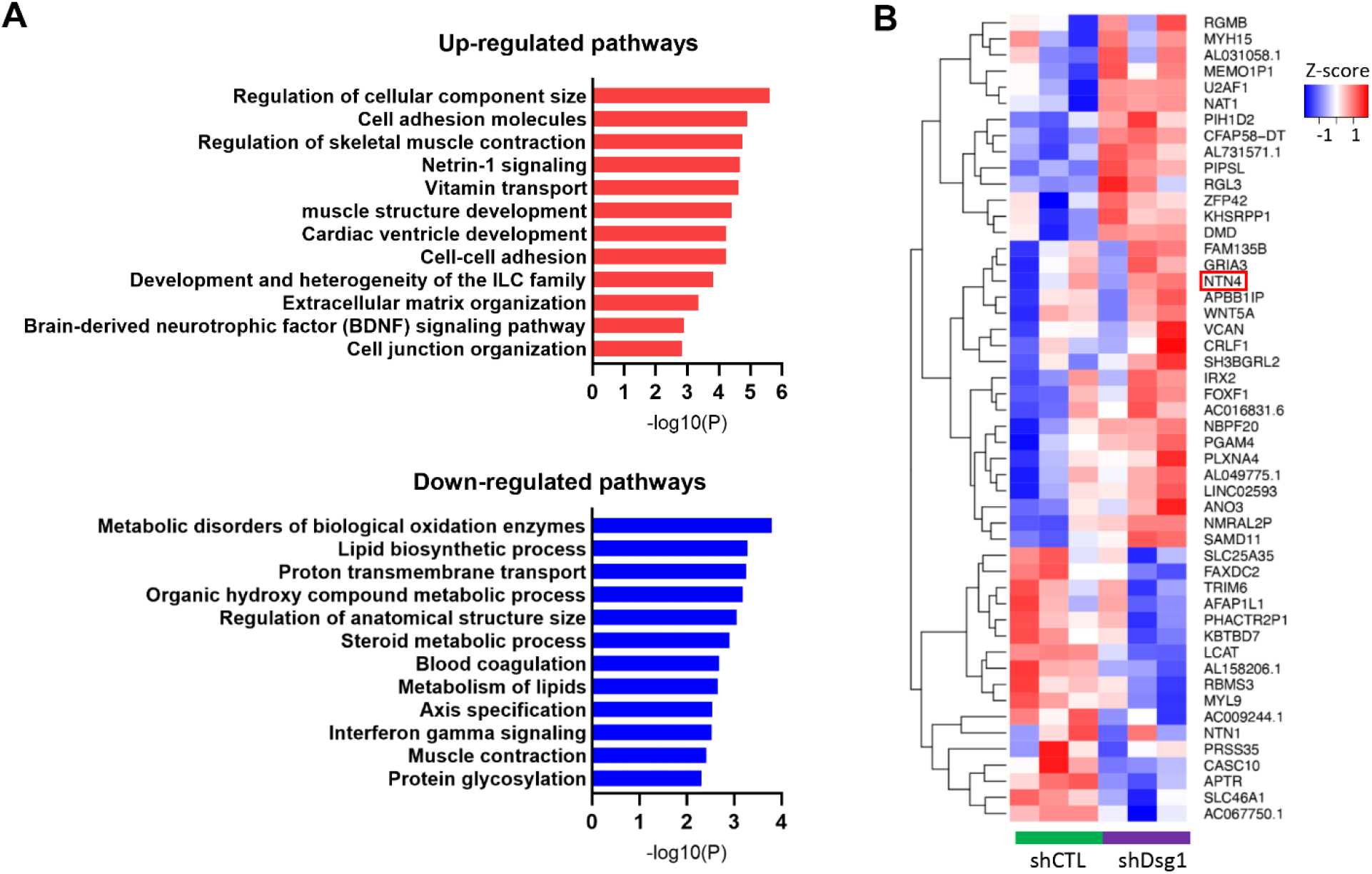
Upregulation of NTN4 gene in BRAF^V600E^ melanocytes treated by Dsg1-deficient CM. **A**, Bar graph of top enriched pathways represented in genes that were significantly up/down-regulated by Dsg1 knockdown CM. **B**, Heatmap with hierarchical clustering revealed the expression of the top 50 differentially expressed genes between BRAF^V600E^ melanocytes treated with control and Dsg1-deficient CM.

To confirm up-regulation of NTN4 gene expression in BRAF^V600E^-transduced cells treated with CM from Dsg1-deficient keratinocytes, qRT-PCR was performed. The results demonstrated a ∼2-fold increase in NTN4 mRNA after treatment with shDsg1 CM (Fig. 4A). In addition, immunofluorescence analysis showed increased staining for Netrin-4 protein upon shDsg1 CM treatment (Fig. 4B). Next, we used shRNAs to knock down NTN4 expression in melanocytes. Of three shRNAs tested, two reduced Netrin-4 expression (both at mRNA level and protein level) (Fig. 4C and 4D). We chose shNTN4_70 for use in subsequent experiments. Melanocytes were first transduced with shNTN4 or shCTL for 2 days, then cells were infected with BRAF^V600E^, followed by treating with control or Dsg1-deficient CM for 3 days. SA-β-Gal assay showed that after knockdown of Netrin-4 expression, Dsg1-deficient CM no longer inhibited BRAF^V600E^-induced senescence (Fig. 4E and 4F), suggesting that the ability of Dsg1-deficient CM to reduce cellular senescence depends on the presence of Netrin-4.

**Figure 4.**
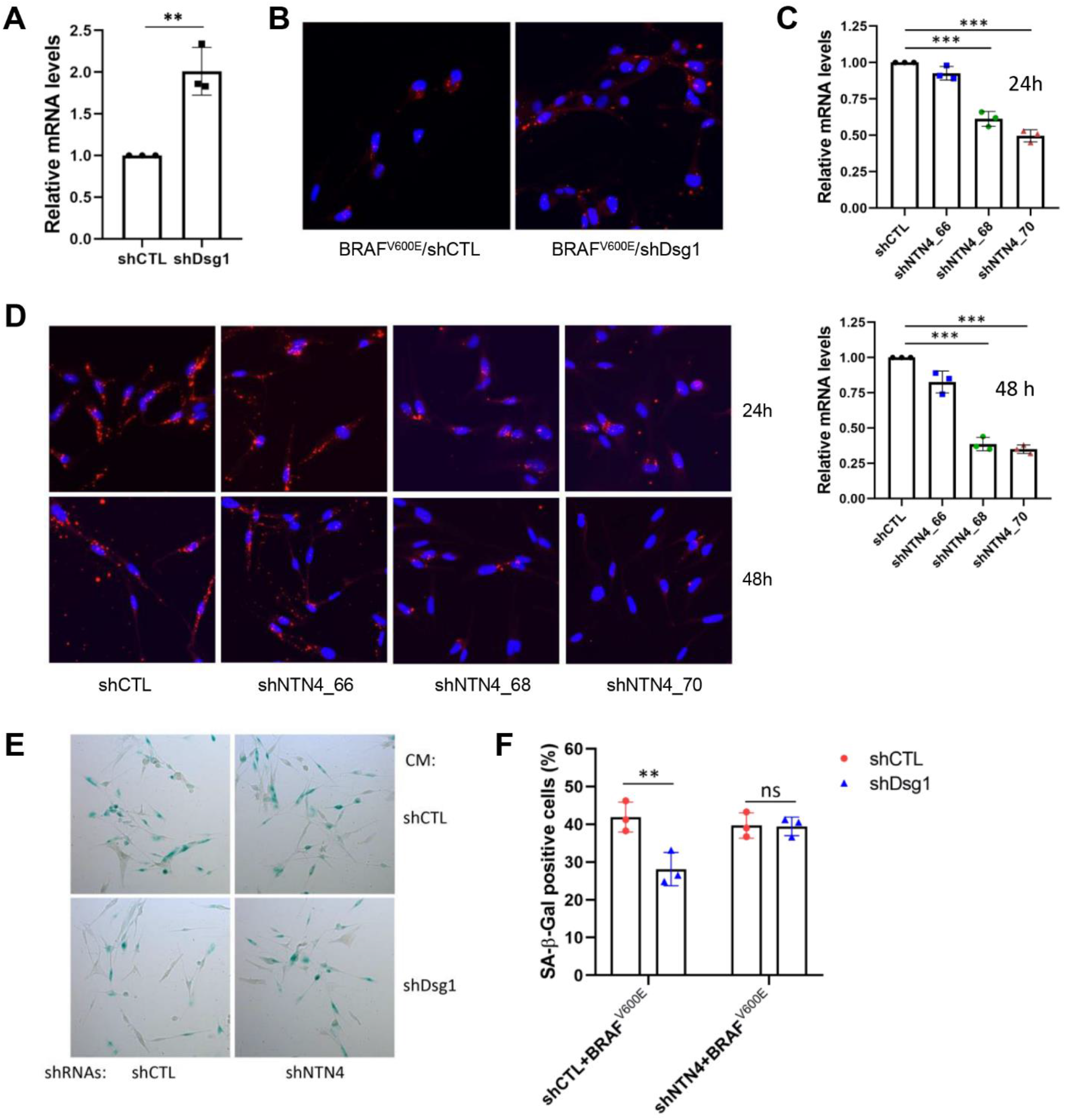
Netrin-4 is involved in Dsg1 loss-induced senescence bypass. **A**, qRT-PCR was performed to confirm the up-regulation of NTN4 expression in BRAF^V600E^-transduced cells treated with Dsg1 knockdown CM. Mean ± SD depicted (n=3), statistical analysis was performed using Student’s t-test (**, p<0.01). **B**, Immunofluorescence analysis confirmed higher expression of Netrin-4 protein upon Dsg1-deficient CM treatment. **C** and **D**, qRT-PCR and immunofluorescence staining were carried out to confirm the knockdown of NTN4 expression (both mRNA and protein) by shRNAs. Results in **C** was reported as Mean ± SD (n=3), statistical analysis was performed using one-way ANOVA with multiple comparisons (***, p<0.001). **E**, Senescence-associated β-galactosidase expression was detected by X-gal staining assay. **F**, Quantification of X-gal staining assay. Mean ± SD depicted (n=3), statistical analysis was performed using two-way ANOVA with multiple comparisons (**, p<0.01).

## Discussion

Cutaneous melanocytic nevi frequently harbor BRAF and NRAS mutations, which drive a burst of cell proliferation [43], but which seldom develop to melanoma due to OIS [10]. However, some malignant melanomas do arise from preexisting nevi, supporting the idea that driver mutation-induced growth arrest is reversible. Further, 80% of melanomas, even those not arising from nevi, harbor activating BRAF mutations, supporting their key role in melanoma development. The acquisition of additional mutations in cell cycle and tumor suppressor genes is thought to be a major path to bypassing growth arrest to promote melanomagenesis. However, less is known about extrinsic, non-genetic factors that could contribute to senescence bypass in the absence of genetic changes. Here we show that paracrine signaling due to loss of Dsg1 in keratinocytes results in transcriptional reprogramming from a more differentiated to less differentiated and more stem-like state, and increased senescence bypass in primary BRAF^V600E^ expressing melanocytes. These data support the idea that loss of keratinocyte Dsg1, possibly occurring in response to acute environmental stress such as UV, could provide an extrinsic stimulus to bypass growth arrest and OIS in nevi to promote the next steps in melanocyte transformation and melanonomagenesis (Fig. 5).

**Figure 5.**
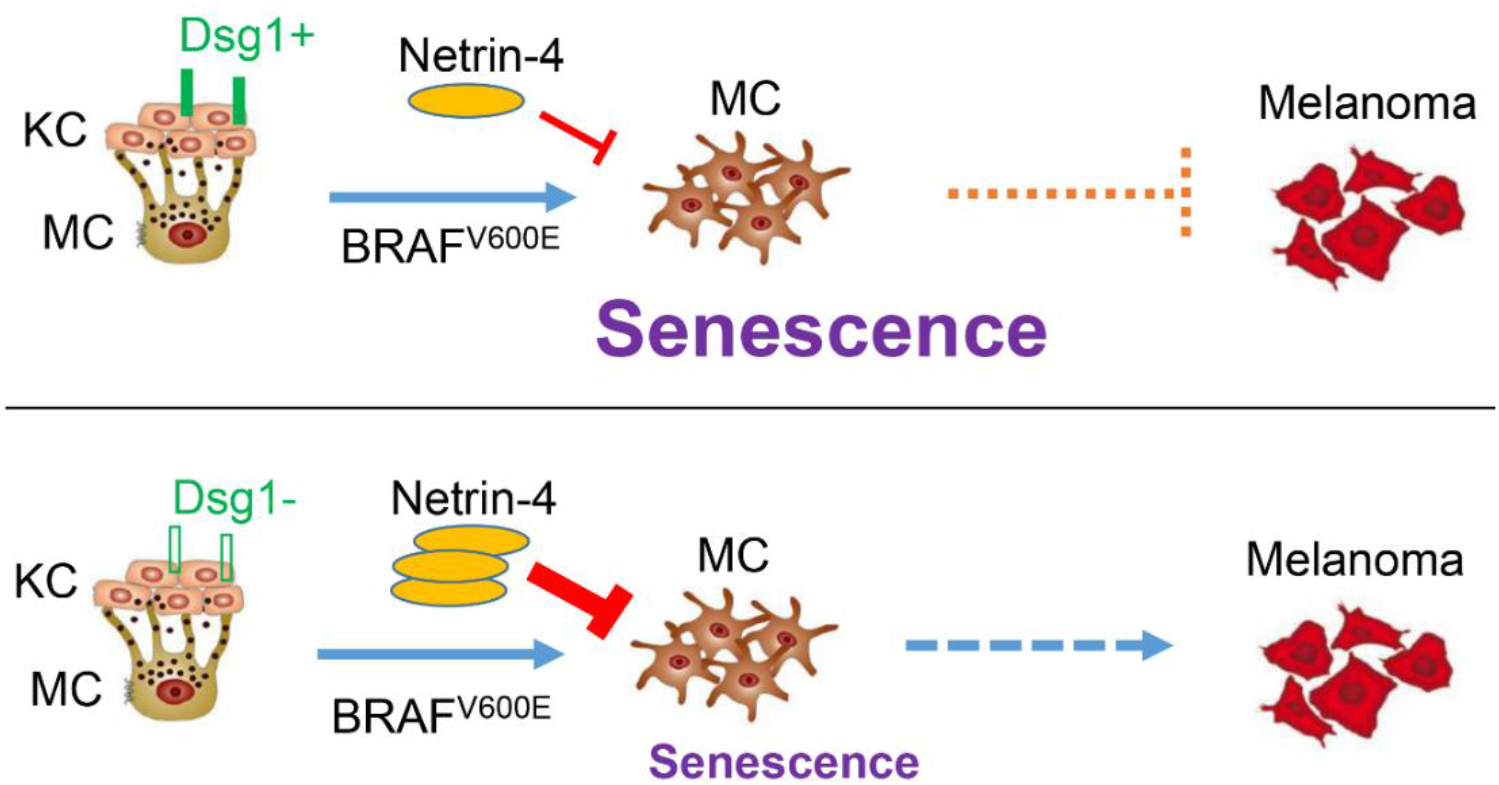
A schematic diagram to indicate the role of Dsg1 in keratinocyte:melanocyte communication and melanomagenesis. The loss of Dsg1 in keratinocytes alters adjacent melanocytes through paracrine signaling and induces Netrin-4 expression, thus promoting cellular senescence bypass.

While it is unclear what components of conditioned media from Dsg1-deficient keratinocytes are responsible for the observed transcriptional and behavioral changes in BRAF^V600E^ melanocytes, a recent report demonstrated that either hyperproliferation or reversible mitotic arrest can occur in human melanocytes depending on whether Tetradecanoylphorbol acetate (TPA) is present in the media [12]. TPA is a protein kinase C (PKC) activator used to substitute for endothelin receptor type B activation as a mitogen in primary melanocyte media. It was shown that in the presence of TPA, BRAF^V600E^ expression inhibited cell proliferation; however, in TPA-free media, BRAF^V600E^ expression induced proliferation. Transcriptional analysis showed that this behavior was accompanied by enrichment of genes associated with cell adhesion and melanocyte progenitor and stem cells. This is reminiscent of the increase in adhesion-related genes and decrease in differentiated gene signatures we observed in BRAF^V600E^ melanocytes treated with shDsg1 media. It is important to note that in the present study, we used a different culture medium tailored to maintain a differentiated pigment-producing phenotype in melanocytes, and constitutive rather than inducible BRAF^V600E^ expression system, both could influence the results.

Among the mechanisms by which extrinsic factors can impact tumor initiation and progression are epigenetic changes, for instance through histone or chromatin modifications and noncoding RNAs [44, 45]. In the study alluded to above, beyond affecting the differentiation status of cells, TPA also induced expression of MIR211-5p, which was required for BRAF^V600E^ induced mitotic failure. Interestingly, MIR211-5p is transcriptionally suppressed by UV exposure [46], a condition under which Dsg1 is also suppressed and which our experimental design models here.

In the present study we identified both shared and distinct changes in transcriptional reprogramming in control and WT BRAF melanocytes compared with BRAF^V600E^ expressing melanocytes. Signatures shared by all cell types were those associated with invasive or neural crest stem cell associated transcripts. Whereas melanocytes originating from the neural crest ultimately come under control of surrounding keratinocytes once they have migrated to the basal layer of epidermis, melanoma cells that have escaped keratinocyte control can resemble multipotent progenitors [47]. Together, these observations suggest that loss of keratinocyte Dsg1 could contribute to a loss of keratinocyte control over melanocytes. They are also in line with our previous observation that even normal primary melanocytes exhibit local spreading (pagetoid behavior) in long term reconstructed epidermal co-cultures with Dsg1-deficient keratinocytes [21], suggesting that this loss of control can occur early in melanocyte transformation.

Whereas the invasive/stem-like signatures were shared across cell types, BRAF^V600E^-transduced melanocytes were unique in exhibiting a switch from pigmented/melanocytic to dedifferentiated gene signatures. In contrast the control and WT BRAF melanocytes showed the opposite; that is, they switched to more differentiated/melanocytic transcriptional programs, despite also exhibiting the neural crest stem-like signatures. This, again, is in line with our data showing that control primary melanocytes exhibited an increase in pigment secretion when treated with shDsg1 media and is consistent with the idea that the temporary decrease in Dsg1 we see in response to UV radiation may stimulate protective responses like melanin synthesis, that are in part mediate by Dsg1 loss [21]. Our new data indicate, however that once melanocytes are expressing the BRAF^V600E^ oncogene, keratinocyte Dsg1 loss induces transcriptional changes associated with de-differentiation, and coupled with the invasive signatures, would drive these cells down a pro-tumorigenic path.

Among the changed genes in BRAF^V600E^-transduced melanocytes linked with adhesion and differentiation status, NTN-4/Netrin-4 stood out to us based on its role in inhibiting senescence in endothelial cells. Netrins are secreted proteins that were initially identified as guidance cues, directing cell and axon migration during neural development. Subsequent findings have demonstrated that netrins can affect the formation of multiple tissues by mediating cell migration, cell-cell interactions and cell-extracellular matrix adhesion [48]. In the context of tumorigenesis, it was previously demonstrated that Netrin-4 is a lymphangiogenic factor contributing to tumor dissemination and represents a potential target to inhibit metastasis [49]. Netrin-4 also promotes neuroblastoma cell survival and migration [50, 51]. Recent studies showed that Netrin-4 inhibits senescence in endothelial cells and protects glioblastoma cells from TMZ-induced senescence [30-32]. Here, we found that Netrin-4 was up-regulated by Dsg1-deficient CM in BRAF^V600E^-transduced melanocytes. Further, knockdown of Netrin-4 expression suppressed the senescence bypass mediated by Dsg1-deficient CM, indicating senescence bypass through loss of Dsg1 depends on the presence of Netrin-4. Previous studies have shown that Netrin-4 is upregulated by EGF/EGFR signaling [31]. In this respect it is interesting to note that baseline NTN-4 expression is significantly increased in shCTL CM-treated BRAF^V600E^ melanocytes, consistent with the well-established elevation of MAPK signaling in these cells compared to WT BRAF melanocytes. Future work will address the extent to which shDsg1 media further elevates EGFR/MAPK signaling in melanocytes to upregulate Netrin-4 transcription.

Taken together, we propose that loss of Dsg1 in keratinocytes alters adjacent melanocytes through paracrine signaling. Whereas control and WT BRAF melanocytes exhibit transcriptional signatures that are on one hand more melanocytic, they share signs of invasive/stem-like features of neural crest progenitor cells. It is important to note that given the known heterogeneity that can exist even in normal melanocyte populations, we cannot know whether the invasive versus more differentiated signatures seen in control and WT BRAF cells come from the same or distinct cell populations. In the case of BRAF^V600E^-transduced melanocytes, while the presence of this oncogene results in a mitotic block, they nevertheless seem predisposed to further steps in melanocyte transformation, once exposed to a Dsg1-deficient environment. Not only did they appear less differentiated and more invasive/stem-like from a transcriptional standpoint, but also a proportion escaped senescence under these conditions. This ability to bypass senescence appeared at least in part to be due to elevation of the secreted laminin-like protein Netrin-4 in BRAF^V600E^ melanocytes. Together, our data support the idea that loss of keratinocyte Dsg1, which could occur through external environmental stress like UV or signals from transforming melanocytes, could provide extrinsic signals to adjacent melanocytes with OIS, helping a proportion escape this mitotic block to promote melanomagenesis.

## Materials and Methods

### Cell Culture

Keratinocytes and melanocytes were isolated from neonatal foreskin provided by the Northwestern University Skin Biology & Diseases Resource-Based Center (SBDRC) as previously described [52]. Keratinocytes were grown in M154 medium supplemented with human keratinocyte growth supplement (Thermo Fisher Scientific), 1,000 x gentamycin/amphotericin B solution (Thermo Fisher Scientific), and 0.07mM CaCl_2_ (low calcium). Melanocytes were cultured in OptiMEM (Thermo Fisher Scientific) containing 5% fetal bovine serum (MilliporeSigma), 10 ng/ml bFGF (ConnStem), 1 ng/ml heparin (MilliporeSigma), 0.1 mM N6, 2’-O-dibutyryladenosine 3:5-cyclic monophosphate (dbcAMP, MilliporeSigma), 0.1 mM 3-isobutyl-1-methyl xanthine (IBMX, MilliporeSigma), and 1% penicillin/streptomycin (Corning). All cells were maintained at 37 °C with 5% CO_2_.

### Viruses generation and transduction

To knock down Dsg1 expression in keratinocytes by retroviruses, LZRS-shDsg1 (shDsg1) and LZRS-NTshRNA (shCTL) were generated as described [21, 53]. Keratinocytes were transduced with retroviral supernatants produced from Phoenix cells as previously described [54].

To express WT BRAF and BRAF^V600E^ in melanocytes, pLV-Puro-CMV_WT_BRAF-3x-FLAG and pLV-Puro-CMV_MUT_BRAF-3xFLAG lentiviral vectors were purchased from VectorBuilder. To knock down NTN4/Netrin-4 expression, GIPZ non-silencing lentiviral shRNA control and GIPZ human NTN4 lentiviral shRNA (3 different clones: V2LHS_60466, V2LHS_60468, and V2LHS_60470) vectors were purchased from Horizon. Lentiviruses were generated by SBDRC Gene Editing Transduction and Nanotechnology Core. For lentiviral transduction, melanocytes were seeded at 30-40% confluence in six-well plates and the next day incubated in lentiviral supernatants with 8 μg/ml polybrene (MilliporeSigma) for 5-7 hours, then the viral supernatants were removed and replaced with growth media.

### Conditioned media preparation and treatment

Conditioned media (CM) from keratinocytes were prepared by following procedure: Cells were plated at 20% confluence in 10-cm dishes, infected with shCTL or shDsg1 retroviral supernatants, grown to 80% confluence, expanded to three 10-cm dishes, and grown to confluence. At this point, cells were switched to high calcium (1.2 mM CaCl_2_) media and continued to culture for 3 days. On day 3, the media were replaced with fresh high calcium media and harvested on day 5. All CM were stored at -80°C in aliquots until use in experiments. To treat melanocytes with CM, normal melanocyte medium was mixed with CM (1:1), added to melanocyte cell culture and replaced with fresh mixed CM media every 2 days.

### Senescence-associated β-galactosidase assay

For the colorimetric analysis of senescence-associated β-galactosidase (SA-β-Gal), melanocytes were plated in two-well slide chambers. Following lentiviral BRAF transduction and 3-day CM treatment, a SA-β-Gal staining kit (Abcam, ab65351) was used according to the manufacturer’s instructions. Senescent cell quantification was performed by counting SA-β-Gal positive (blue) cells in random images taken using a bright-field microscope. An average of 200 cells/sample were counted.

For the fluorescence analysis of SA-β-Gal, melanocytes were plated in 6-well plate. Following the same treatment as described above, a CellEvent Senescence Green Flow Cytometry Assay Kit (Thermo Fisher Scientific, C10841) was used per the manufacturer’s instructions. Samples were analyzed on a flow cytometer (BD LSRFortessa X-20 Cell Analyzer) using a 488 nm laser. An average of 10,000 cells/sample were counted.

### Western blot

Whole cell lysates were collected from samples in urea-SDS buffer (8 M urea,1% SDS, 60 mM Tris (pH 6.8), 10% glycerol, and 5% ß-mercaptoethanol) after PBS wash, and sheared with a syringe and a 25g needle. Protein concentrations were determined by the BCA assay (Thermo Fisher Scientific). Lysates were separated by SDS-PAGE and transferred to nitrocellulose membrane. The membrane was blocked with 5% non-fat dry milk for 1 hour at room temperature and followed by incubation with the primary antibody solution overnight at 4 °C or 1 hour at room temperature. After a series of PBS washes, the membrane was incubated with HRP-linked secondary antibodies for 1 hour at room temperature. Protein bands were imaged using LI-COR Odyssey FC (LI-COR Biosciences). Densitometric analysis was performed on blots using LI-COR Image studio software. The following primary antibodies were used: BRAF (#9433), p44/42 MAPK (#9107), P-p44/42 MAPK (#4370), and p16 (#80772) from Cell Signaling Technology, BRAF^V600E^ (ab200535) from Abcam, Dsg1 (#32-6000) from Thermo Fisher Scientific, GAPDH (G9545) from MilliporeSigma, and Tubulin (12G10) from Developmental Studies Hybridoma Bank. Secondary antibodies for immunoblotting were goat anti-mouse and goat anti-rabbit peroxidase (SeraCare Life Sciences).

### EdU assay

Melanocytes were seeded in 6-well plate, transduced with lentiviral WT BRAF or BRAF^V600E^ and followed by treatment of CM for 3 days. EdU was added into culture media at a 10 μM final concentration 24 hours before samples were harvested. The Click-iT Plus EdU Flow Cytometry Assay Kit (Thermo Fisher Scientific, C10632) was used according to the manufacturer’s instructions. EdU incorporation into DNA was analyzed by flow cytometry.

### Flow cytometry analysis of cell death

Dead Cell Apoptosis Kit with Alexa Fluor 488 Annexin V and PI for Flow Cytometry (Thermo Fisher Scientific, V13241) was applied to determine cell death after BRAF transduction and CM treatment described as above. Cells were washed with cold PBS and resuspended with annexin V binding buffer, 5 µl Alexa Fluor 488 annexin V and 1 µl 100 μg/mL PI working solution were added to 100 µl of cell suspension. Cells were then incubated in the dark at room temperature for 15 min and another 400 µl of binding buffer was added. Data were collected on flow cytometer and were analyzed with FlowJo.

### RNA sequencing

Human normal melanocytes, melanocytes transduced with either WT BRAF or BRAF^V600E^ were cultured with CM from control or Dsg1 knockdown keratinocytes for 3 days. Total RNA was extracted from cells using the Quick-RNA miniprep kit (Zymo Research) per the manufacturer’s instructions and sent to Novogene for mRNA library preparation and transcriptome sequencing. Data analyses were performed in R using Bioconductor libraries and R statistical packages. Differential expression analysis and batch correction were performed using DESeq2. Gene set enrichment analyses were conducted using GSEA. Gene annotations and pathway enrichment analysis were carried out by Metascape [55].

### qRT-PCR

Total RNA was extracted from cells using the Quick-RNA miniprep kit (Zymo Research) as described above. cDNA was synthesized by using the Superscript III First Strand Synthesis Kit (Thermo Fisher Scientific) from 1-2 µg of total RNA. Quantitative PCR was performed on the QuantStudio 3 instrument (Thermo Fisher Scientific), using SYBR Green PCR master mix (Thermo Fisher Scientific). Relative mRNA levels were calculated through the ΔΔCT method and normalized to endogenous GAPDH expression. Gene-specific primers for human NTN4 (F: 5’-GTACTTTGCGACTAACTGCTCC-3’, R: 5’-TCCAGTGCATGGAAAAGGACT-3’) and GAPDH (F: 5’-CATGAGAAGTATGACAACAGCCT-3’, R: 5’-AGTCCTTCCACGATACCAAAGT-3’).

### Immunofluorescence

Melanocytes were plated into 2-well chamber slide (Nunc Lab-Tek II), either infected with lentiviral BRAF^V600E^ and treated with CM or transduced with lentiviral NTN4 shRNAs. Cells were then washed with PBS, fixed by 4% paraformaldehyde for 15 minutes, followed by permeabilization with 0.2% Triton X-100 in PBS for 5 minutes, and blocked by 1% BSA/2% normal goat serum in PBS for 1 hour at room temperature. Primary antibody Netrin-4 (R&D, AF1254) was diluted in the blocking buffer (1:100) and incubated overnight at 4 °C. The following day, cells were washed 3 times with PBS and incubated with an Alexa-568 conjugated secondary antibodies (ThermoFisher Scientific) for 1 hour at 37 °C. DAPI was then added to the cells and incubated for 2 minutes for nuclear staining. After mounting with ProLong Gold Antifade Mountant (ThermoFisher Scientific), Immunofluorescence images were captured by fluorescence microscopy (Carl Zeiss, AxioVison Z1 system).

### Statistical analysis

Statistical analysis was performed using GraphPad Prism 9.0 software. All experiments were performed at least three times. Measured data were represented as the mean ± SD. One-way or two-way analysis of variance (ANOVA) with multiple comparisons or Student’s t-test was applied to compare quantitative data. P-values for each analysis are marked on figures, and the level of statistical significance was defined as * p<0.05, ** p<0.01, and *** p<0.001.

## Acknowledgements

This work was supported by NIH R01 CA228196, AR041836, AR043380, and the J.L. Mayberry Endowment to KJG. Research was also supported by NIH P30 AR075049 awarded to Northwestern University Skin Biology & Diseases Resource-Based Center, with special thanks to the GET iN and STEM cores.

