## Supplemental Figures for "Crosstalk in skin: Loss of desmoglein 1 in keratinocytes inhibits BRAF^V600E^-induced cellular senescence in human melanocytes"

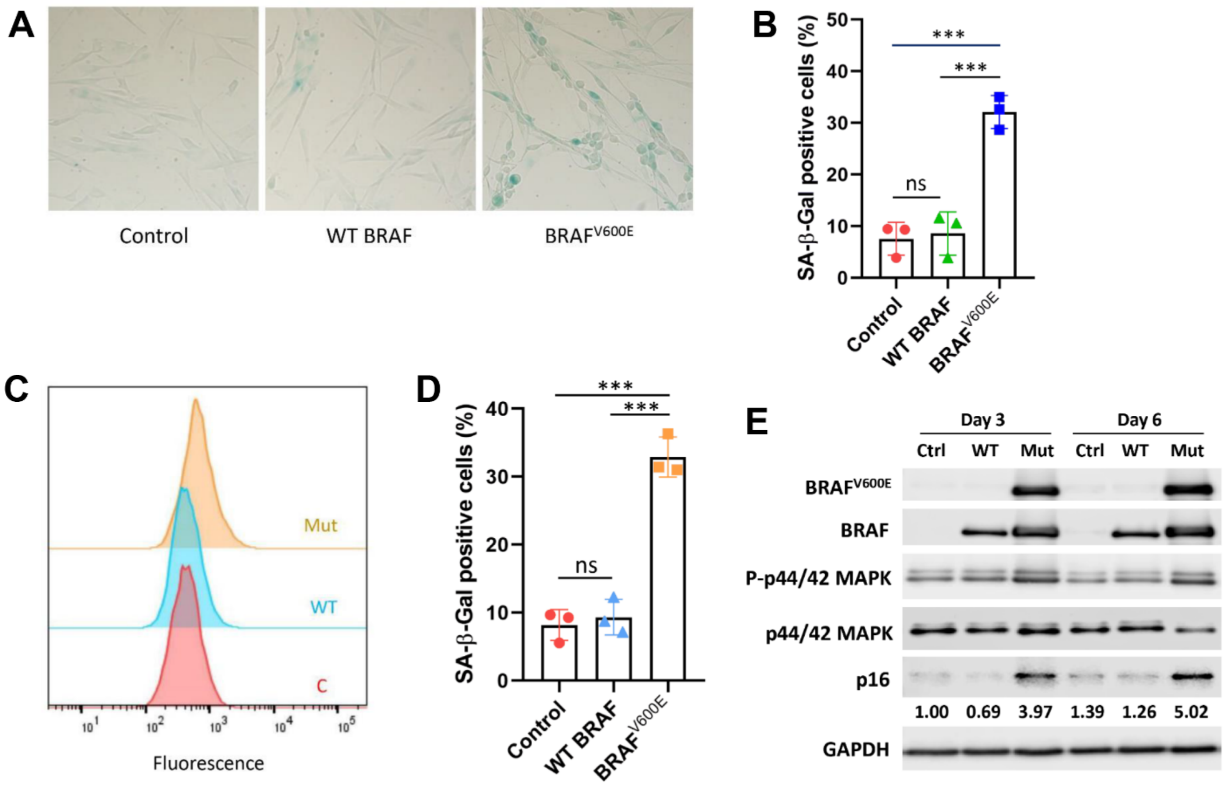


**Figure S1. BRAF$^{V600E}$ induces cellular senescence in human melanocytes.** Melanocytes were transduced with WT BRAF or BRAF$^{V600E}$ lentivirus and cultured for 3 or 6 days before they were harvested. **A,** Senescence-associated β-galactosidase (SA-β-Gal) expression was detected by X-gal staining assay. **B,** Quantification of X-gal staining assay. **C,** SA-β-Gal expression was detected by flow cytometry assay. Right shift of the peak indicated the increase in senescence. **D,** Quantification of senescence flow cytometry assay. **E,** Expression of the indicated proteins was determined by Western blot, a significant increase in p16 levels was observed 3 days after lentivirus transduction and lasted for 6 days. Results were reported as Mean ± SD (n=3), statistical analysis was performed using one-way ANOVA with multiple comparisons (***, $p<0.001$).

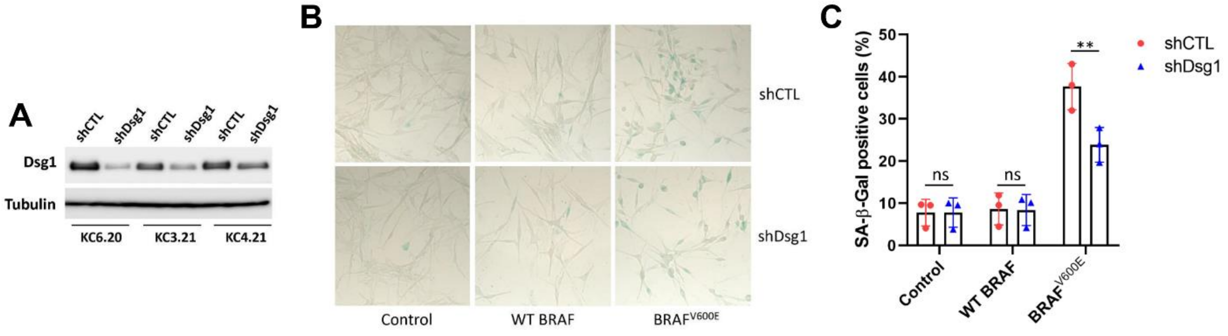


**Figure S2. Conditioned media generation and X-gal staining to detect senescence after CM treatment. A,** Western blot to demonstrate the knockdown of Dsg1 expression in keratinocytes (3 keratinocyte clones from 3 different foreskin were tested). **B,** Senescence-associated β-galactosidase expression was detected by X-gal staining assay. **C,** Quantification of X-gal staining assay. Mean ± SD depicted (n=3), statistical analysis was performed using two-way ANOVA with multiple comparisons (**, p<0.01).
